# Amyloidogenic regions in beta-strands II and III modulate the aggregation and toxicity of SOD1 in living cells

**DOI:** 10.1101/2023.10.18.562627

**Authors:** Luke McAlary, Jeremy R Nan, Clay Shyu, Mine Sher, Steven S. Plotkin, Neil R. Cashman

**Author notes:** Correspondence to: Luke McAlary Neil Cashman.

## Abstract

Mutations in the protein superoxide dismutase-1 (SOD1) promote its misfolding and aggregation, ultimately causing familial forms of the debilitating neurodegenerative disease amyotrophic lateral sclerosis (ALS). Currently, over 220 (mostly missense) ALS-causing mutations in the SOD1 protein have been identified throughout the primary sequence, indicating that common structural features responsible for aggregation and toxicity may be present. Here, we used *in silico* tools to predict amyloidogenic regions in the ALS-associated SOD1-G85R mutant, finding 7 regions spread throughout the protein structure. We found that the introduction of proline residues into β-strands II (I18P) or III (I35P) reduced the aggregation propensity and toxicity of SOD1-G85R in living cells, significantly more so than proline mutations in other amyloidogenic regions. The I18P and I35P mutations also reduced the capability of SOD1-G85R to template onto previously formed non-proline mutant SOD1 aggregates as measured by fluorescence recovery after photobleaching. Finally, we found that, while the I18P and I35P mutants are less structurally stable than SOD1-G85R, the proline mutants are less aggregation-prone during proteasome inhibition, and less toxic overall. Our research highlights the importance of a previously underappreciated SOD1 amyloidogenic region in β-strand II (^15^QGIINF^20^) to the aggregation and toxicity of SOD1 in ALS mutants, and suggests that β-strands II and III may be good targets for the development of SOD1-associated ALS therapies.

## Introduction

Protein aggregation is associated with numerous human diseases, in particular those involving neurodegeneration such as Alzheimer’s disease, Parkinson’s disease, and amyotrophic lateral sclerosis (ALS) (1). Although the major pathological hallmark of ALS is the deposition of the RNA/DNA-binding protein TAR-DNA-binding protein of 43 kDa (TDP-43) into insoluble inclusions (2), the first protein genetically linked to ALS was Cu/Zn superoxide dismutase (SOD1) (3). Mutations in SOD1 are currently thought to be responsible for ∼20% of familial ALS (FALS) cases and ∼1% of sporadic cases (4). At this time, over 200 ALS-associated mutations are reported to occur within SOD1 (5), although the mechanisms by which they cause ALS remain elusive. The most well-accepted hypothesis is that the misfolding and subsequent aggregation of SOD1, as a consequence of mutation, is somehow toxic to motor neurons (6).

SOD1 is a homodimeric superoxide scavenging enzyme in which each monomer contains an intrasubunit disulfide bond and binds both a zinc ion and a copper ion (7). These post-translational modifications afford substantial stability to the protein, making it resistant to thermal and chemical denaturation *in vitro* (8, 9). Considering the high stability of the SOD1 native conformation, extensive research has been carried out examining the effect that ALS-associated mutations have on the folding stability of the protein under various conditions (10–12). There appears to be a general trend of mutations to cause decreased folding stability, where some mutations lead to decreased metal-binding and/or increased susceptibility of reduction of the intrasubunit disulfide bond, or even dimer dissociation. Interestingly, many SOD1 mutants, when folded properly, retain wild-type-like enzymatic activity (7), suggesting that mutations are likely affecting the protein folding landscape early in SOD1’s maturation/folding pathway (13). Regardless of the exact effect a mutation has on SOD1 folding stability, they all result in the deposition of SOD1 into cytoplasmic inclusions in SOD1-FALS patient motor neurons.

An interesting feature of SOD1-FALS is that SOD1 misfolding and aggregation can be propagated intra- and intercellularly *in vitro* and *in vivo* in a prion-like manner (14–17). For example, lipofection of cultured cells expressing SOD1-G85R tagged with green fluorescent protein (GFP) with homogenates from spinal cord tissue of SOD1-FALS patients leads to increased SOD1-G85R-GFP inclusion formation in recipient cultures (18). Also, injection of spinal cord homogenates from symptomatic SOD1 mice into non-symptomatic SOD1 mice can induce ALS-like phenotypes and pathology (16). Furthermore, it was recently shown that injection of SOD1-FALS human patient spinal cord homogenates into transgenic SOD1 mice was capable of inducing ALS pathology and symptoms (19). Together, these data suggest that SOD1 can act in a prion-like manner to spread misfolding and aggregation from cell-to-cell.

The ability to propagate in a prion-like manner suggests that there is an underlying substructure to SOD1 aggregates that may be responsible for this. Some evidence suggests that SOD1 inclusions in ALS patients can be composed of small granule-coated fibrils (20), whereas other evidence suggests that the proteinaceous inclusions are amorphous (21). *In vitro* aggregation assays of recombinant SOD1 protein have shown that even the highly stable wild-type (WT) protein can form amyloid-like fibrils when reduced and metal-free (12, 22, 23). Furthermore, several studies using binary epitope mapping of the insoluble protein fraction from the spinal cords of transgenic SOD1 mice identified specific regions of the protein to be consistently inaccessible to antibodies in the insoluble material (17, 19, 23). This binary epitope mapping was also capable of distinguishing between two separate strains of SOD1 aggregates, characterized by differing antibody-inaccessible regions of the protein (17). These regions encompass some *in silico* predicted and experimentally verified amyloidogenic regions of SOD1 (24–26), supporting a role for amyloid-like aggregation in SOD1-FALS.

Although these amyloidogenic regions of SOD1 were identified (24), only one was examined in cell-based models of SOD1 toxicity (27). A small synthetic peptide of the predicted amyloidogenic region ^28^PVKVWGSIKGL^38^, corresponding to beta-strand III, was found to form corkscrew-shaped oligomers composed of out of register beta-sheets (27). These oligomers were toxic when exogenously added to cultured primary neurons, and a tryptophan substitution at Gly33 could alleviate toxicity of the peptide and full length non-native SOD1 oligomers (27). This region of the protein is particularly interesting within the context of SOD1 aggregation and prion-like propagation as (i) it contains few ALS-associated mutations compared to the rest of the amino acid sequence, (ii) it nevertheless has high sequence divergence from murine SOD1 (murine SOD1 does not co-aggregate with human SOD1 or become misfolded in murine models (15, 28)), and (iii) contains the sole tryptophan in SOD1 which plays a key role in SOD1 aggregation and prion-like propagation (15, 29, 30). Although it is clear that the sequence region ^28^PVKVWGSIKGL^38^ is important to SOD1-FALS aggregation, what is still unknown is what effect the other amyloidogenic regions have on SOD1 aggregation and toxicity in the context of full-length SOD1 when expressed inside mammalian cell lines, as it is in human disease. Indeed, addition of exogenous protein to cultured cells can be toxic through different pathways compared to when the protein is expressed within cells (31). We set out to address these questions in detail using well-established cell culture models of SOD1 aggregation and toxicity (12, 18, 29). By introducing proline substitutions into each amyloidogenic region of a SOD1-G85R-AcGFP construct, we mapped the contribution of each region to the intracellular aggregation and toxicity of SOD1-G85R. We specifically identified that two N-terminal amyloidogenic regions (^15^QGIINF^20^, and ^30^KVWGSIKGL^38^) are the most important for cellular aggregation and cytotoxicity. We further establish that reduction of the amyloidogenicity of mutant SOD1 with 7 proline substitutions significantly reduces its aggregation propensity and toxicity in cultured motor neuron-like cells, even though this particular mutant is highly destabilized and unfolded as measured by its susceptibility to proteolytic digestion. Lastly, we determine that the amyloidogenicity of ^15^QGIINF^20^, and ^30^KVWGSIKGL^38^ regions impact the ubiquitin proteasome-mediated degradation of misfolded SOD1. Overall, our data suggest that the N-terminus, in particular beta-strands II and III, of SOD1 are responsible for its conformational conversion into toxic aggregates.

## Results

### Cu/Zn superoxide dismutase contains several regions predicted to be amyloidogenic

Although SOD1 is not generally considered to be a classical amyloidogenic protein based upon histopathological evidence (lack of Thioflavin-S and congo-red binding in patient tissue), it is still capable of forming amyloid fibrils *in vitro* (12, 22, 23, 25, 29). Indeed, previous work predicted and confirmed several amyloidogenic regions within SOD1(24) using the 3D-profile method (32). Inspired by these findings, we hypothesized that potentially more amyloidogenic regions could be identified by combining the outputs of several *in silico* sequence-based amyloidogenic prediction tools and previously published work. Previous work using recombinant SOD1 protein identified a core region of SOD1 fibrils corresponding to residues 1-63 (25), whereas other work, also using recombinant SOD1, identified several core regions including residues 1-40, 90-120, and 130-153 (26). Also, whilst not strictly identifying amyloid, binary epitope mapping of insoluble SOD1, from model human mutant SOD1 expressing mice, identified buried regions of different SOD1 aggregate strains (Strain A: residues 4-39 and 80-127, Strain B: 4-57) (17).

To this end, we loaded the human SOD1-G85R sequence into several computational prediction tools (see Material and Methods for details) in order to predict amyloidogenic regions within the protein. On the basis of the combined outputs, we predicted 7 amyloidogenic regions within the SOD1 sequence, which, as expected, correlated with beta-strand structures within the protein **(Figure 1 A and B)**. Previous work only predicted 4 amyloidogenic regions in SOD1 (^14^VQGIINFE^21^, ^30^KVWGSIKGL^38^, ^101^DSVISLS^107^, and ^147^GVIGIAQ15^3^) (24), however, only a single predictor (3D-profile method) was utilized in that case. Our predictions overlap with these regions; In addition, regions not previously predicted included ^4^AVCLVKG^10^, ^85^RNVTAD^90^ (in the G85R mutant), and ^113^IGRTLVVH^120^, which correspond to strands I, V, and VII respectively **(Figure 1 A and B)**. Note these regions show distinct non-overlap and anticorrelation with predicted solvent exposed misfolding-specific epitopes of SOD1 (Peng et al, DOI: 10.1021/acs.jpcb.8b07680). We then introduced proline substitutions into each of these regions to identify the residue location that would decrease predicted amyloidogenicity the most, identifying A4, I18, I35, A95, I104, I112, T116, and I149 **(Supp. Fig. 1)**. Interestingly, proline substitutions at I18, I35, I104, and I149 were all predicted previously to block fibril formation, but *in vitro* purified protein aggregation assays showed that only I104P and I149P were capable of blocking fibril formation to a certain degree (24). Mutation of 7 prolines in all amyloid regions (7P mutant) resulted in a substantial reduction in overall predicted amyloidogenicity **(Supp. Fig. 2)**

**Figure 1.**
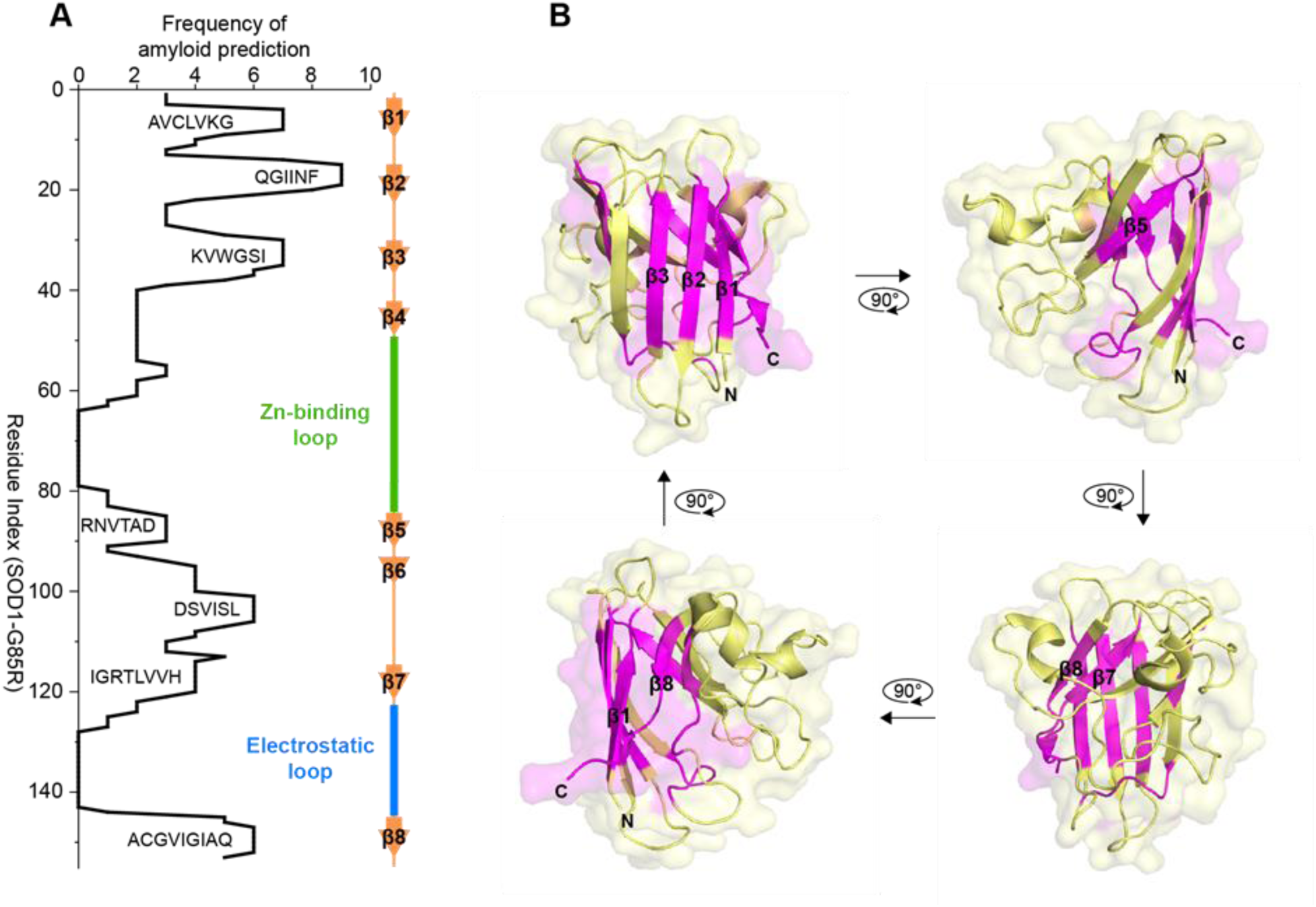
Predicted amyloidogenic regions in SOD1-G85R. **(A)** The frequency of each residue in the SOD1-G85R sequence to be predicted as amyloidogenic from several prediction tools and previous literature (see methods). **(B)** The location of the amyloidogenic regions (magenta) in the context of the SOD1 monomeric structure (PDB: 1HL5(33)).

### SOD1-FALS mutant aggregation in cultured cells is primarily mediated by β-strands II and III

Following the identification of the amyloidogenic regions in SOD1 and the amino acids to substitute proline, we generated mammalian expression vectors to express SOD1-G85R C-terminally tagged with AcGFP (34) and made single proline substitutions at our identified sites. We generated A4P, I18P, I35P, A95P, I112P, T116P, I149P, and 7P (all sites substituted with proline) mutants on the SOD1-G85R-AcGFP backbone. SOD1 tagged with fluorescent protein at the C-terminus is a common model for studying the effects of mutations on the cellular aggregation of SOD1 (12, 18, 29, 35–37). We have previously used the SOD1-G85R-AcGFP vector in several studies, where it has responded to drug treatment (29) and prion-like seeding by SOD1-FALS patient tissue (18).

In order to determine the effect the proline substitutions would have on the formation of cellular inclusions, we transiently transfected these vectors into U2OS cells and analysed the cells using fluorescence microscopy **(Figure 2A)**. We observed that U2OS cells transfected with either AcGFP, SOD1 wild-type (WT), I18P, I35P, or 7P showed diffuse GFP signal within the majority of cells when visualized via fluorescent microscopy **(Figure 2A)** and signal was present at levels greater than that of G85R **(Supp Fig 3)**. In contrast, G85R, A4P, A95P, I104P, I112P, T116P, and I149P all resulted in inclusions within the cytoplasm of at least some cells **(Figure 2A)**. To gain a more quantitative understanding of the effect the proline substitutions were having on the formation of inclusions in cells, we used an unbiased machine-learning-based approach (38–40) to identify cells with or without inclusions from fluorescent microscopy images. We observed, under the experimental conditions used here, that cells expressing either I18P, I35P, or 7P did not significantly differ in inclusion formation compared to WT expressing cells, with the percentage of transfected cells containing inclusions ranging from 2-4% **(Figure 2B)**. Cells expressing either G85R, A4P, A95P, I104P, I112P, T116P, or I149P formed significantly more inclusions than WT expressing cells, indicating that these amyloidogenic regions are less important to the aggregation of SOD1 in cells than those encompassing I18 or I35, an interesting result considering previous work showed no effect of I18P or I35P substitutions on SOD1 fibril formation *in vitro* (24). Comparison of the inclusion formation of all mutants to G85R expressing cells showed that WT, A4P, I18P, I35P, A95P, and 7P formed significantly less inclusions, however, A4P and A95P had values that were closer to G85R than the other mutants with ∼7% and 10% of cells containing inclusions respectively **(Figure 2B)**.

**Figure 2.**
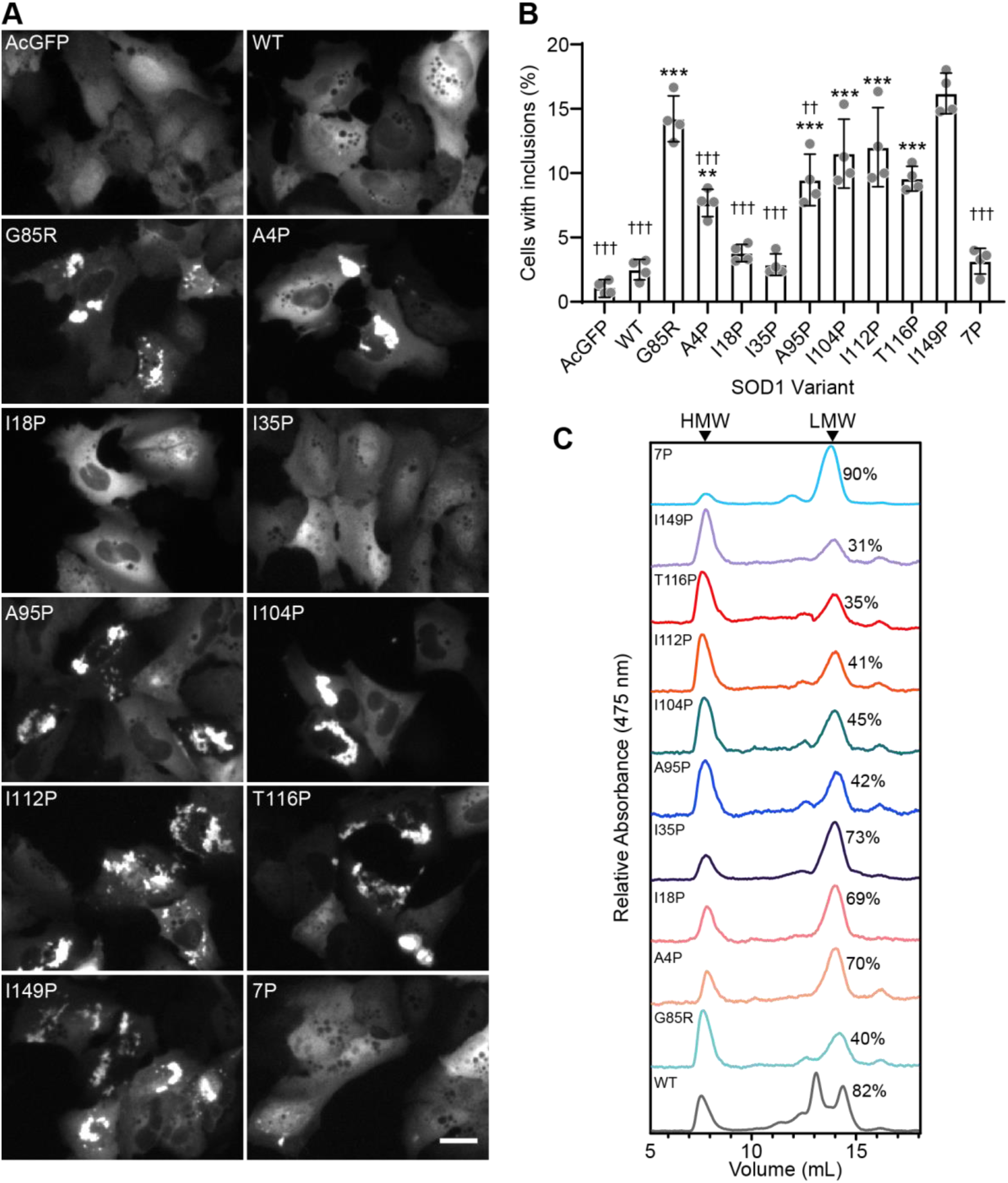
Inclusion formation and aggregation of SOD1-G85R is reduced by disruption of N-terminal amyloidogenic regions but not those in the C-terminus. **(A)** Representative images of U2OS cells transfected with plasmids expressing AcGFP-tagged human SOD1 variants showing aggregates (bright puncta). Scale bar = 25 µm. **(B)** Quantification of the extent of inclusion formation observed in U2OS cells 48 h post-transfection. **(C)** Analytical size-exclusion chromatography measuring absorbance at 475 nm of lysates from cells expressing SOD1 variants showing high molecular weight (HMW) species and low molecular weight (LMW) species. Numbers indicate the percentage of signal (area under the curve) that was in the LMW fraction. (*** signify comparisons with SOD1-WT; ††† signify comparison to G85R). Significance was determined with a one-way ANOVA with Tukey’s post-test (*** or ††† *p < 0.001, \*\** or †† *p < 0.05, \** or † *p < 0.01,* ns = not significant).

Additionally, we examined the level of SOD1-AcGFP oligomers formed in cells by performing analytical size-exclusion chromatography (SEC) on cell lysates, measuring the absorbance of AcGFP at 475 nm **(Figure 2C)**. Cells were lysed and insoluble material was pelleted via centrifugation to ensure that only soluble particles were analysed. We observed that there were two major peaks for all variants, with the exception of SOD1-WT, which showed three peaks. The two low-molecular weight (LMW) peaks present for SOD1-WT are monomer and dimer (eluting at ∼14.5 and 13 mL respectively). For other variants, the single LMW peak is likely unfolded monomers as has been observed previously for destabilised SOD1 analysed by gel-filtration chromatography (12). The high-molecular weight (HMW) peak for each variant was observed at ∼7.5 mL and is the void volume of the column used (Superdex 75; **Figure 2C)**, and is potentially SOD1-AcGFP oligomers and/or SOD1 bound to other proteins. By measuring the area under the curve for each peak, we were able to determine the percentage of SOD1-AcGFP present as LMW or HMW species for each variant, finding that the variants with the greatest proportion of LMW were A4P, I18P, I35P, and 7P. These variants also resulted in the lowest number of transfected cells with inclusions along with WT **(Figure 2B)**.

### Proline mutations in amyloidogenic segments β-strands II or III prevent templated aggregation of SOD1 in living cells

Previous work has shown that mutant SOD1 expressed in cultured cells is capable of being seeded to aggregate using preformed seed from recombinant SOD1 aggregates (41), tg SOD1 mouse spinal cord homogenates (42), or SOD1-FALS patient spinal cord homogenates (18) in a prion-like manner. Considering this, we wished to examine how our proline substituted SOD1-G85R-AcGFP constructs would respond to the presence of aggregated SOD1 in cells: Would it template through other aggregation-prone regions or not? In order to investigate this, we co-transfected our cells with a SOD1-G85R-TdTomato construct and either AcGFP, SOD1-WT-AcGFP, or SOD1-G85R-AcGFP proline mutant constructs, where SOD1-G85R-TdTomato aggregates and act as a template for AcGFP-tagged constructs. We have previously used this system to investigate seeded aggregation in cells by quantifying the relative fluorescent intensities of each fluorescent protein (either AcGFP or TdTomato) within inclusions as a function of its total fluorescence in the cell (29).

Co-transfection of U2OS cells with SOD1-AcGFP and SOD1-G85R-TdTomato vectors resulted in the formation of protein inclusions within the cell cytoplasm that were positive for both GFP and TdTomato signal, although to different extents dependent on the specific proline substitution or lack thereof **(Figure 3A)**. Quantification of this, by determining the ratio of fluorescent signal in an inclusion and total cytoplasm, showed that both AcGFP alone and SOD1-WT-AcGFP were less abundant in inclusions than SOD1-G85R-TdTomato (*P* < 0.0001) **(Figure 3B)**. Conversely, SOD1-G85R-AcGFP was slightly more likely to partition to inclusions than SOD1-G85R-TdTomato, akin to our previous work involving Trp32 substitutions in SOD1 (29). The ratio of proline mutants I18P, I35P, and 7P in SOD1-G85R-TdTomato inclusions did not significantly differ from that of SOD1-WT-AcGFP or AcGFP, indicating a resistance to templated aggregation in this assay. In comparison, cells transfected with A95P, I104P, I112P, T116P, and I149P showed similar levels in SOD1-G85R-TdTomato inclusions for these proline mutants, indicating a similar aggregation propensity.

**Figure 3.**
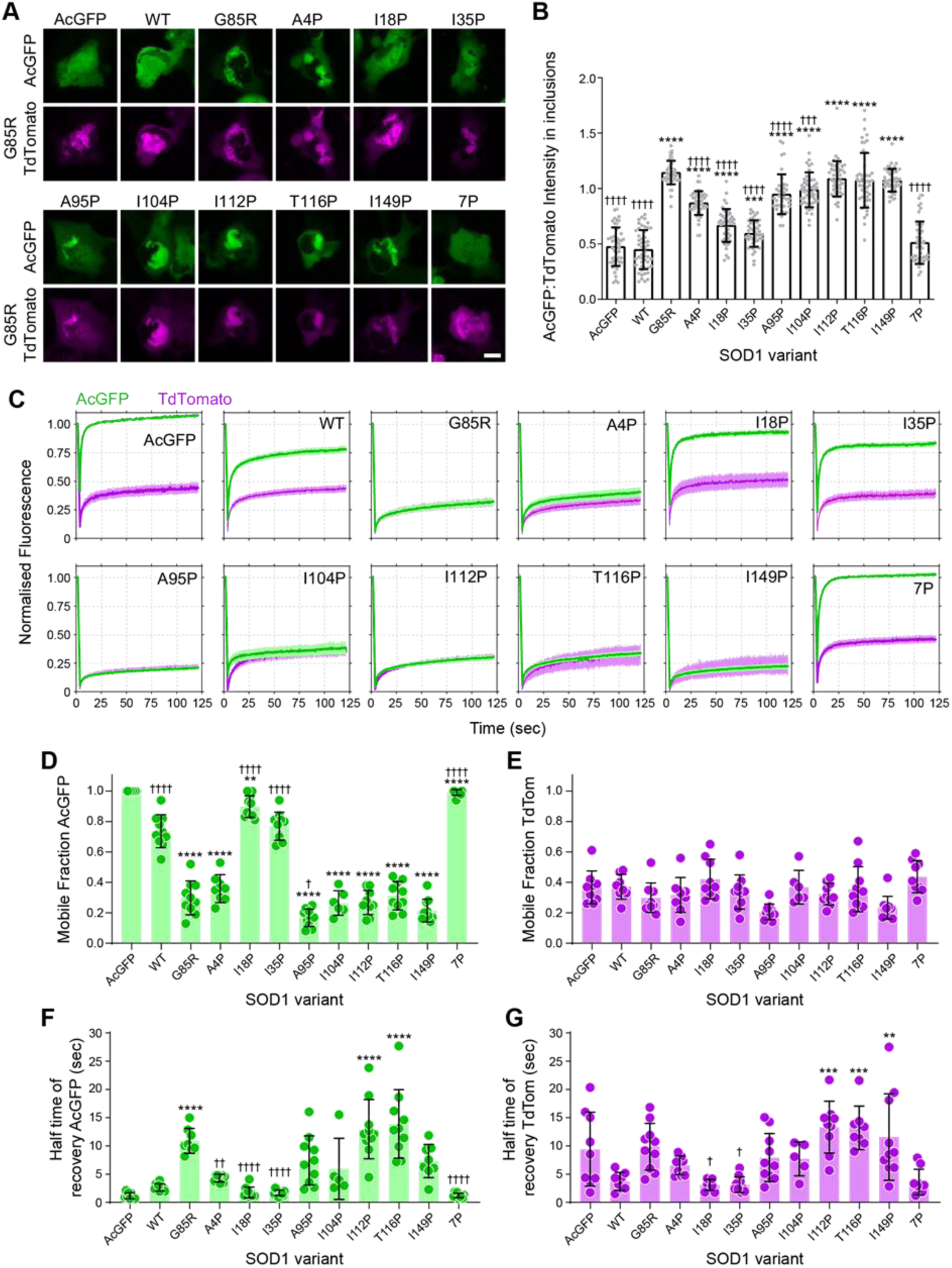
I18P and I35P variants resist co-aggregation with SOD1-G85R-Tdtomato inclusions. **(A)** Representative images of U2OS co-transfected with SOD1-AcGFP constructs (green) and SOD1-G85R-TdTomato (magenta). Scale bar = 10 µm. **(B)** Quantification of the ratio of AcGFP signal in TdTomato inclusions comparative to free AcGFP in living cells. **(C)** Fluorescence Recovery After Photobleaching (FRAP) of inclusions in cells co-expressing AcGFP-tagged SOD1 variants (green) and SOD1-G85R-TdTomato (magenta). **(D)** Quantification of the mobile fraction of AcGFP-tagged SOD1 variants from FRAP curves. **(E)** Quantification of the mobile fraction of SOD1-G85R-Tdtomato from FRAP curves. **(F)** Quantification of the halftime of recovery of AcGFP-tagged SOD1 variants from FRAP curves. **(G)** Quantification of the halftime of recovery for SOD1-G85R-TdTomato from FRAP curves.

To gain more insight into the dynamics of SOD1 within inclusions in our cell model, we used fluorescence recovery after photobleaching (FRAP). FRAP exploits the bleaching of fluorescent molecules to measure the movement of non-bleached fluorescent molecules into a region-of-interest (ROI) that has been bleached (43). This allows for the determination of the fraction of fluorescent molecules within an ROI that are mobile, and also the time it takes for unbleached molecules to diffuse into a bleached ROI. By co-transfecting cells with both SOD1-AcGFP proline variants and SOD1-G85R-TdTomato we were able to use the TdTomato signal to determine inclusion ROI’s for bleaching. We reasoned that FRAP would allow us to more accurately determine the fraction of SOD1 within an inclusion that is actually part of the aggregate and not simply free soluble SOD1 that is in the same ROI.

We observed, under our FRAP parameters, that both AcGFP and TdTomato were detectable and bleachable with a 488 nm argon laser (**Supp Fig 4**). Owing to this, we measured the mobile fraction and recovery time of both AcGFP and TdTomato in our assay. When bleaching AcGFP without SOD1 conjugated to it the mobile fraction was calculated to be close to 1.0, indicating AcGFP was not part of SOD1-G85R-Tdtomato inclusions (**Figure 3 C**). However, SOD1-WT-AcGFP had a mobile fraction of 0.74 ± 0.11 **(Fig 3 C and D**), indicating that roughly a quarter of the SOD1-WT-AcGFP was immobile in the inclusions. Interestingly, I18P and 7P variants were both significantly more mobile than WT in inclusions with mobile fractions of 0.90 ± 0.07 (*P* = 0.011) and 0.99 ± 0.02 (*P* < 0.0001) respectively **(Fig 3 C and D**), indicating that these variants were even more resistant to template directed aggregation than WT SOD1. The mobile fraction of SOD-G85R-AcGFP was significantly lower than WT, I18P, I35P, and 7P variants at 0.30 ± 0.11 (*P* < 0.0001) but not significantly different from A4P, I104P, T112P, I116P, and I149P, suggesting that these proline substitutions did not affect template directed aggregation. Meanwhile, the A95P variant had a significantly lower mobile fraction than SOD1-G85R-AcGFP at 0.17 ± 0.06 (*P* = 0.0198), suggesting that this substitution increased susceptibility to template directed aggregation. Measuring the mobile fractions for the SOD1-G85R-TdTomato that was co-transfected with the AcGFP-tagged SOD1 variants showed no significant differences between G85R-AcGFP transfected cells and other variants **(Fig 3 E**).

We next measured the half-time of recovery for each AcGFP variant to determine if there were differences in the diffusion speed of the mobile fraction within inclusions. The half-time to recovery for the AcGFP-tagged variants was rapid and similar to WT for those proline mutants that were less aggregation prone (A4P, I18P, I35P, 7P) and some aggregation-prone mutants (A95P, I149P) indicating some variance in the mobility of SOD1-AcGFP in inclusions. Interestingly, the half-time to recovery for SOD1-G85R-TdTomato also showed similar variance, where values correlated with those of the SOD1-AcGFP variants **(Supp Fig. 5)**, indicating that the diffusion speed of SOD1-G85R-TdTomato in inclusions was affected by the SOD1-AcGFP variant it was co-expressed with.

### Intracellular Aggregation propensity correlates with SOD1 proline variant toxicity in cultured motor neuron-like cells

Previous work has indicated that the levels of SOD1 inclusion formation in cells may correlate with cell toxicity. Other work has suggested that soluble species are responsible for cellular toxicity. Considering that our I18P, I35P, and 7P variants are all highly soluble but still contain the disease-causing G85R mutation, we wanted to determine relative toxicity of these variants in a motor neuron-like cell model. To this end, we transfected the SOD1 variants into NSC-34 cells and performed live-cell microscopy to observe the formation of protein inclusions and cell counts across time **(Figure 4A)**.

**Figure 4.**
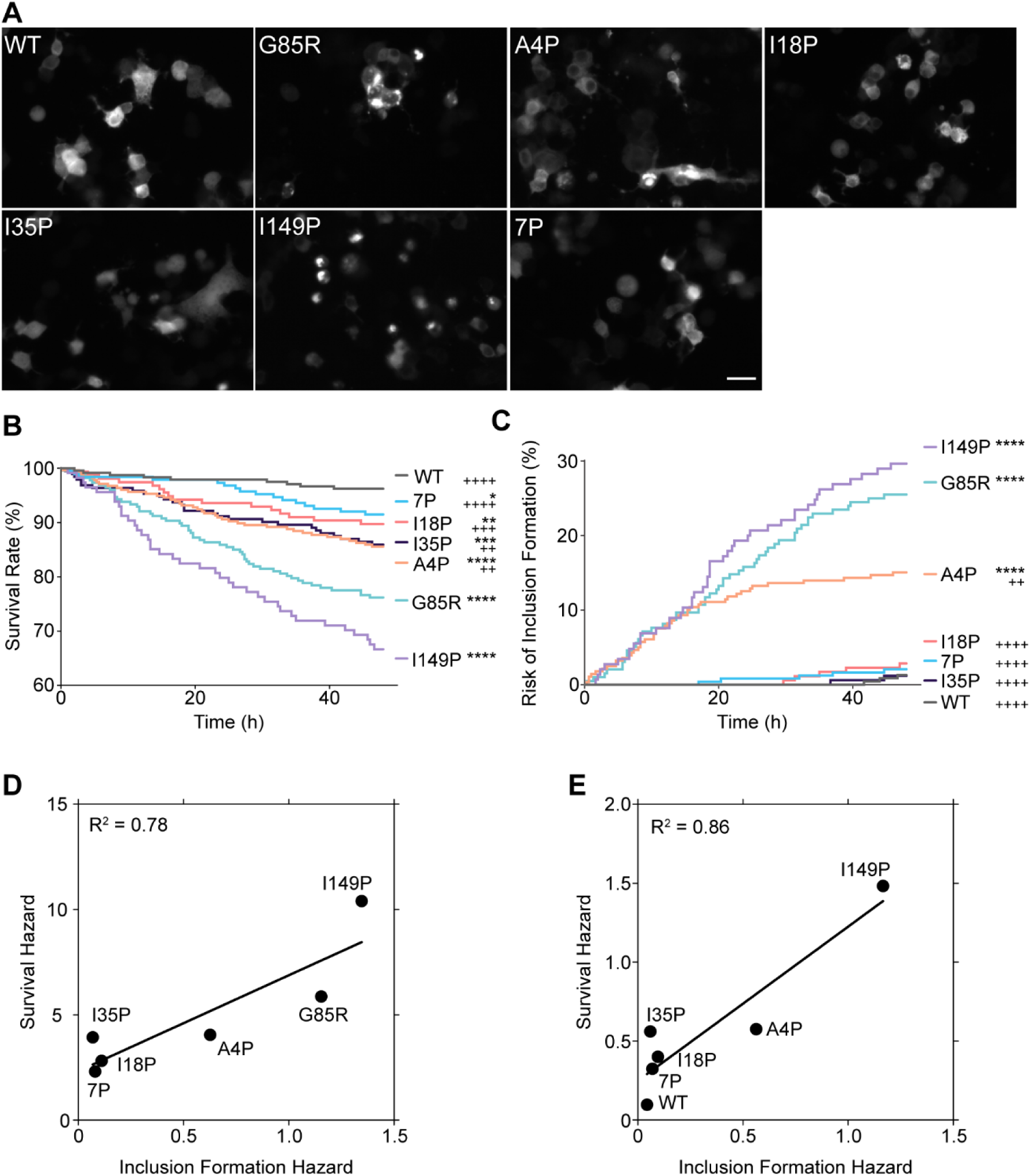
I18P and I35P variants partially detoxify SOD1-G85R in living motor neuron-like cells. **(A)** Representative images of NSC-34 cells expressing SOD1-AcGFP variants. **(B)** Survival analysis of variants over a 48 h period where events are when a cell died. **(C)** The risk of inclusion formation was determined by determining the time that a cell first presented with an inclusion visually. **(D)** Linear regression of survival hazard against inclusion formation hazard from comparisons of variants to SOD1-WT. **(E)** Linear regression of survival hazard against inclusion formation hazard from comparisons of variants to SOD1-G85R. Survival and hazard curves were generated from 3 separate experimental repeats. Comparisons between curves were made using Log-rank (Mantel-Cox) test. Hazard ratios were determined by log-rank method. * = comparisons made against SOD1-WT and ^+^ = comparisons made against SOD1-G85R. **** = P < 0.0001, *** = P < 0.001, ** = P < 0.005, and * = P < 0.01, with symbols meaning the same for ^+^ comparisons.

We applied Kaplan-Meier analysis to both cell death **(Figure 4B)** and inclusion formation **(Figure 4C)** to determine the risk of these events occurring for each SOD1 variant examined (WT, G85R, A4P, I18P, I35P, I149P, and 7P). For cell survival, we found that SOD1 proline variants were all still significantly more likely to die than SOD1-WT in this model, even including the 7P variant **(Figure 4 B)**, suggesting ablation of amyloidogenicity does not fully detoxify misfolded SOD1 in living cells. Comparisons made to SOD1-G85R showed that all of the proline variants, with the exception of SOD1-I149P, were more significantly more likely to survive to varying degrees, suggesting some effect at reducing toxicity of SOD1-G85R. We next monitored the formation of inclusions in our live-cell image stacks to determine the relative risk of inclusion formation in this model **(Figure 4 C)**. We observed that only A4P, G85R, and I149P variants resulted in significantly more inclusions in NSC-34 cells when comparisons were made against WT. On the other hand, all variants except I149P formed significantly less inclusions in comparison to G85R. These data match those seen above for U2OS cells **(Figure 2)**.

We next determined that there were appreciable positive correlations between survival hazard and the hazard of inclusion formation for comparisons made to WT **(Figure 4 D)** and G85R **(Figure 4 E)**, with R^2^ values being 0.78 and 0.86 respectively. This indicates that the level of SOD1 inclusion formation in this model is correlated with the death of the NSC-34 cells.

### The 7P variant is structurally destabilized but more resistant to intracellular degradation than SOD1-G85R

Previous work has shown that the half-life of SOD1 in various models is mutation-dependent, where the WT is long-lived and ALS-associated mutants are shorter-lived (44–47). The overarching hypothesis has been that mutations in SOD1 destabilize the protein and promote its degradation, however, there has been no in-depth study to investigate the interplay between stability and degradation. Our previous observation that the hypothetically destabilized 7P variant was highly expressed in cells **(Supp Fig 3)** contradicts the expectation that destabilized forms of SOD1 are degraded only due to instability. Therefore, we decided to investigate the turnover of SOD1 in the context of our SOD1 proline variants. We first set out to determine the relative stabilities of our SOD1 variants using limited proteolysis via both thermolysin and proteinase-K-based assays, and to examine intracellular degradation using cycloheximide to block protein synthesis.

Proteinase-K (PK) is a serine protease that is typically used to detect the conversion of proteins into a prion-like state due to prion-like particles being resistant to proteolysis (48). SOD1, however, is resistant to proteolysis by proteinase-K if it is in its highly stable native conformation (29), meaning that proteinase-K can be used to interrogate the stability of SOD1 variants that reduce the protein’s stability. Incubation of cell lysates containing SOD1-AcGFP variants with proteinase-K revealed that the WT was the most resistant to PK (∼10% decrease in protein levels after 6 h incubation) and 7P and A95P were the most vulnerable, with no detectable protein after 2.5 min **(Supp Fig 6)**. We observed that SOD1-G85R and most proline variants were highly vulnerable to PK digestion, suggesting the population of SOD1 inhabiting the native conformation was low for all variants except for SOD1-WT **(Supp Fig 6)**. To complement this time-based proteolysis assay, we performed fast parallel proteolysis, in which protein samples are differentially heated in the presence of the protease thermolysin so that a temperature-dependent stability can be determined (49). From this, we observed again that WT was the most stable of the variants (∼50% degraded at 65 °C) and 7P was the most unstable, being degraded ∼80% at 4 °C **(Figure 5 A and B)**. SOD1-G85R was more stable than the proline variants, being approximately 50% degraded at 46 °C **(Figure 5 A and B)**. These proteolysis results indicated that even single proline substitutions resulted in substantial destabilisation of the SOD1-G85R structure.

**Figure 5.**
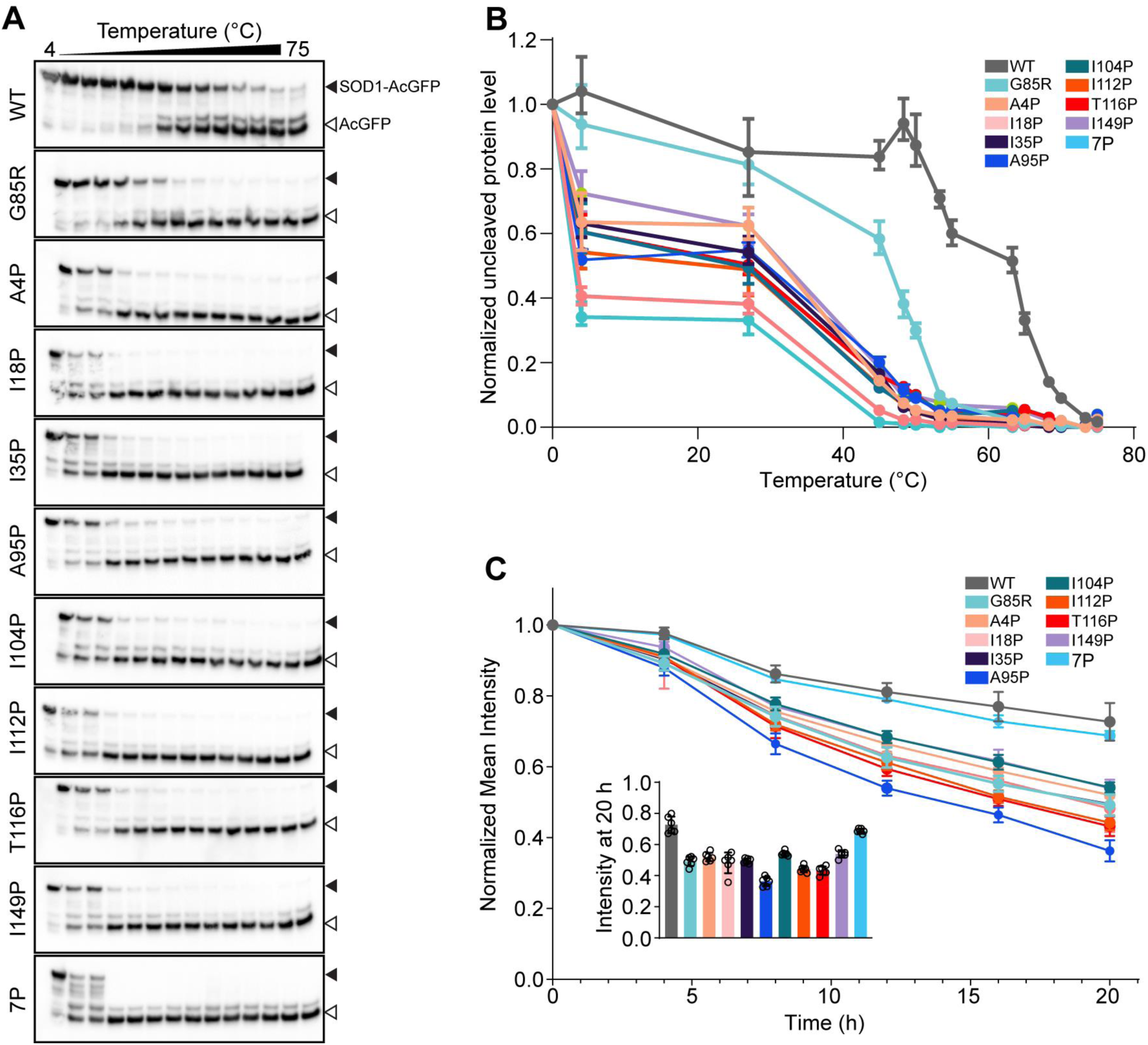
The 7P variant of SOD1-G85R is highly structurally unstable but degraded at a similar rate to SOD1 wild-type in living cells. **(A)** Representative immunoblots from fast parallel proteolysis (FASTpp) experiments. Top band is intact SOD1-AcGFP and bottom band is thermolysin degraded SOD1-AcGFP. **(B)** Quantification of the ratio of intact to cleaved protein across temperatures from FastPP experiments. **(C)** Fluorescence intensity normalised to time 0 h of AcGFP over time in cells expressing SOD1 variants after treatment with cycloheximide to inhibit protein translation. Inset is the data at time = 20 h for easier inspection of differences.

We next sought to understand if the instability of our mutants, as measured via proteolysis, would translate into more rapid turnover in living cells. To do this, we blocked protein synthesis using cycloheximide (CHX) and then monitored GFP fluorescence intensity levels using microscopy over a 20 h time period (imaged every 4 h). Interestingly, we observed that both 7P and WT GFP intensity levels were similar across the CHX treatment period, both decreasing to ∼ 70% at 20 h post-treatment. The other variants decreased much more rapidly and to a greater extent, being at roughly 55% or lower of starting protein levels at 20 h post-treatment. These data indicate that SOD1-WT and a 7P variant are less likely to be degraded in cells than our other proline variants.

### I18P and I35P variants are not substrates of the ubiquitin proteasome, unlike other SOD1 variants

Mutant SOD1 is known to cause dysfunction to the ubiquitin proteasome system (UPS), resulting in cellular toxicity (50–52). Indeed, the inclusions found in animal models of SOD1-fALS and in patients are polyubiquitinated, suggesting a role for ubiquitin signalling and/or the UPS in ALS pathobiology (53). Currently, the exact mechanism through which mutant SOD1 may elicit UPS dysfunction remains unclear, although there are suggestions that an accumulation of polyubiquitinated ALS-associated proteins reduce the pool of free-ubiquitin in cells, leading to aberrant ubiquitin signalling and recycling (50). Considering our results thus far suggested minimal aggregation of some of our variants, we were interested to examine how our unstable but aggregation resistant variants (I18P, I35P, 7P) of SOD1-G85R interacted with the UPS.

We first examined the effect of the proteasome inhibitor MG132 on the inclusion formation of our proline variants since proteasome inhibition is known to increase the levels of aggregated mutant SOD1 (54). Here, a dose-dependent increase in the percentage of cells containing inclusions was observed for all proline-substituted SOD1 variants examined, except WT, I18P, I35P, and 7P which showed no significant increase even at the maximum MG132 concentration tested (10 µM) compared to AcGFP alone **(Figure 6 A and B; Supp Fig 7),** indicating that these variants are not substrates of the UPS like other SOD1 mutants. We further supported this finding by immunoblotting whole and soluble fractions of MG132 treated cells expressing our SOD1 variants, which showed that significantly more SOD1-AcGFP was present in the triton-soluble fraction for WT, A4P, I18P, I35P, and 7P when compared to G85R **(Figure 6 C and D)**. Finally, we determined the relative fraction of each SOD1 variant that was accumulating into ubiquitin-positive inclusions via co-transfection of the variants with mCherry-Ubiquitin (mCh-Ub). By measuring the ratio of SOD1-AcGFP co-localised with mCh-Ub-positive inclusions via confocal microscopy, we determined that all variants except I18P, I35P, and 7P were sequestering to mCh-Ub-positive inclusions to a greater extent than SOD1-WT. Together, these data show that I18P and I35P mutations prevent the sequestration of G85R into ubiquitin-positive inclusions, indicating that aggregation through amyloid segments around I18 and I35 potentiate the formation of toxic SOD1 forms in the cell.

**Figure 6.**
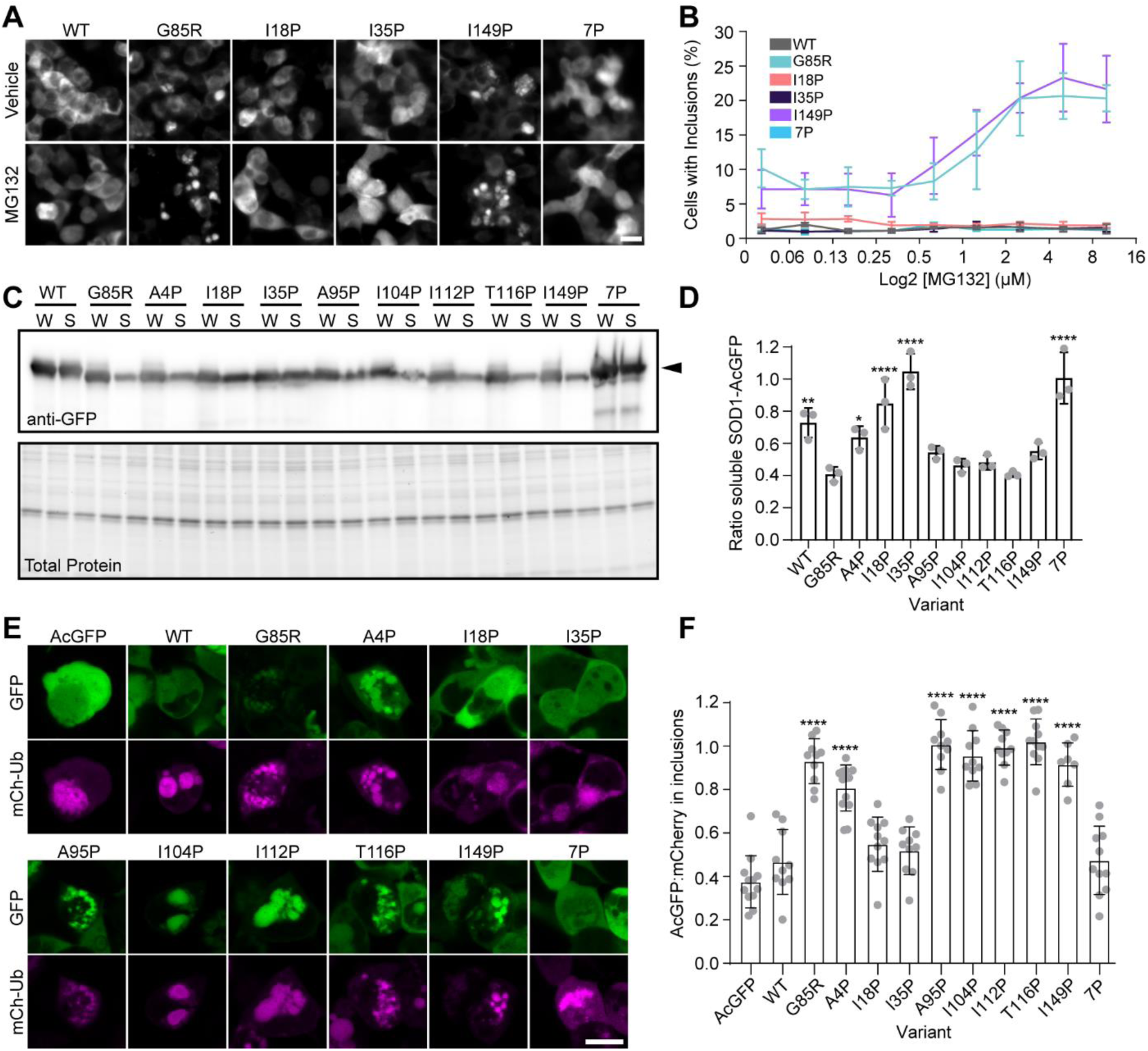
I18P, I35P, and 7P variants of SOD1-G85R are not substrates of the ubiquitin proteasome system (UPS) and do not affect UPS function. **(A)** Micrographs of SOD1-AcGFP variants treated with or without MG132 (10 µM). **(B)** Plot of select SOD1-AcGFP variants (WT, G85R, I18P, I35P, I149P, and 7P) showing the effect of increasing concentrations of MG132 on inclusion formation in cells. **(C)** Immunoblot of lysates from cells expressing SOD1-AcGFP variants treated with 10 µM MG132 (W = whole cell lysate, S = Soluble fraction). **(D)** Quantification of immunoblots from (C), showing that some proline variants do not become insoluble when cells are treated with MG132. **(E)** Micrographs of cells co-transfected with SOD1-AcGFP variants and mCherry-Ubiquitin (mCh-Ub). **(F)** Quantification of the ratio of AcGFP signal in mCh-Ub inclusions comparative to free AcGFP and free mCh-Ub in transfected cells. Error bars represent SD of the mean from at least 3 biological replicates. (**** signify comparisons with SOD1-WT; Significance was determined with a one-way ANOVA with Tukey’s post-test (**** or ††† p < 0.0001, *** < 0.001, ** < 0.01, * p < 0.05). Scale bars are 20 µm.

## Discussion

The formation of protein inclusions and aggregates is considered to be the primary hallmark pathology of most neurodegenerative diseases, and is potentially a causally related process in disease (55). In SOD1-associated ALS, inclusions containing mutant SOD1 are found throughout degenerating tissues, indicating that SOD1 misfolding and aggregation plays a central role in disease pathogenesis (56). Whilst debate exists as to the exact underlying structure of SOD1 in histopathologically-relevant inclusions and whether it is amorphous or amyloid, there is strong evidence to suggest that some sequence regions play a preferential role in SOD1 aggregation (17, 19, 23, 24, 28, 57). Here, using a combination of cell culture, fluorescence microscopy, and immunoblotting, we have determined that predicted fibril forming segments of SOD1 protein contribute differentially to the aggregation and inclusion formation of SOD1 in living cells. Specifically, we identify the amyloidogenic sequences, ^14^VQGIINFE^21^, and ^30^KVWGSIKGL^38^ (corresponding to β-strands II and III respectively) as being the sequence segments that contribute most to the intracellular aggregation of SOD1 in living cells.

It is well-documented that purified recombinant SOD1 can form amyloid fibrils *in vitro* under physiologically relevant conditions, where mutations and post-translational modification can augment both the aggregation kinetics and fibril structures (12, 22, 23, 25, 26, 29, 42). Furthermore, small sequence segments of SOD1 have been identified as having high amyloidogenic propensity, potentially being responsible for SOD1 amyloid formation (24, 27). However, evidence for an amyloid state of SOD1 in animals and humans is not definitive (57). Direct observation of amyloid structures composed of SOD1 *in vivo* is lacking, however, there is strong evidence to suggest structural polymorphs of insoluble SOD1 species are present in animal and cell models of SOD1-associated ALS (16, 17, 19, 23, 28, 36). Interestingly, previous work using binary epitope mapping determined that the most common antibody-inaccessible epitopes of insoluble transgenic human SOD1 in mice corresponded to amino acids 4-20 and 20-39 (23, 28), the former of which encompasses the predicted amyloidogenic regions ^4^AVCLVKG^10^ and ^14^VQGIINFE^21^, whereas the latter encompasses the predicted region ^30^KVWGSIKGL^38^.

Furthermore, *in vitro* and *in silico* force spectroscopy experiments on full-length SOD1 suggest that the N-terminal β-strands form a stable core in the SOD1 unfolding trajectory, supporting the notion that this region could be the common nucleus for pathogenic aggregation (58, 59). In conjunction with the above, our data showing that I18P and I35P variants aggregate less and are less likely to co-aggregate with aggregation-prone SOD1 support this and suggest that β-strands II and III primarily control the initiation and growth of SOD1 aggregates in living cells potentially through formation of a amyloidogenic core.

In regards to β-strand III, there is strong evidence to suggest β-III as the most important segment related to SOD1 aggregation and toxicity. It has been previously determined that the sole tryptophan residue of SOD1 (Trp32; found in segment ^30^KVWGSIKGL^38^) plays an important role in the aggregation of SOD1 and can be targeted with small molecules to reduce aggregation and toxicity in cells and animals (29, 30, 60). Other previous work identified that the ^28^PVKVWGSIKGL^38^ segment forms toxic oligomers when it alone is synthetically produced and aggregated (27). Interestingly, murine SOD1 is resistant to intermolecular prion-like seeding of SOD1 (15) and does not co-aggregate with transgenic human SOD1 in murine models of SOD1-associated ALS (28), a possible consequence of this region exhibiting the least sequence similarity between mice and humans across the SOD1 sequence (29). Indeed, the density of known ALS-associated mutations in and around β-III is much lower than that of the rest of the protein (57). However, canine degenerative myelopathy is associated with mutations in SOD1 (61), where β-III similarity compared to humans is low (29), indicating other regions or factors are at play too.

We did not expect the β-strand II amyloidogenic region to contribute as much as it did to SOD1 aggregation. In fact, in some of our assays, the I18P mutant was more protective than I35P, suggesting a substantial contribution of the ^14^VQGIINFE^21^ segment to SOD1 aggregation. Also of interest, I18P or I35P substitutions both significantly reduced SOD1 aggregation, oligomerization, and toxicity to almost the low levels of 7P and WT. This suggests that β-II and β-III are both required to cause pathological aggregation of SOD1. Within the context of previous work, the segment ^14^VQGIINFE^21^ was identified as amyloidogenic and formed fibrils as a synthetic peptide. A proline substitution in this region in full-length recombinant SOD1 still resulted in fibril formation *in vitro*, supposedly via other amyloidogenic segments in SOD1 (24), which resulted in this region being delegated as unimportant. Previous work has determined that SOD1 interacts with endoplasmic reticulum-associated protein degradation factor Derlin-1 through a Derlin-1 binding site (^5^VCVLKGDGPVQGII^18^) which includes some of ^14^VQGIINFE^21^ (62–64). A potential mechanism by which our I18P variant could be reducing SOD1 mutant aggregation and toxicity is by disrupting the interaction between SOD1 and Derlin-1, which has been shown previously to lower toxicity in cultured cells and SOD1 animal models (63, 64), although we note that I18 is on the very edge of the Derlin-1 binding site. Furthermore, disruption of Derlin-1 binding does not explain the effects we observed of the I35P substitution to reduce toxicity and aggregation as it is distal from the Derlin-1 binding site, suggesting that other mechanisms of mutation-induced toxicity, likely linked to aggregation, are occurring. Indeed, not all mutants of SOD1 have been confirmed to expose this Derlin-1 binding site (65), suggesting that a Derlin-1-SOD1 interaction is just one of many potential pathological consequences of SOD1 mutation, although it remains one of high interest.

Our data here show that the ^14^VQGIINFE^21^ region is important when SOD1 is expressed and forms aggregates within cells, suggesting a role for the cellular environment in the formation of SOD1 oligomeric and aggregate structural polymorphs. Indeed, the advent of cryo-electron microscopy has shown that *ex situ* amyloid fibrils from multiple diseases have substantially different structures in comparison to those formed *in vitro* from recombinant proteins. A recent cryo-electron microscopy-derived structure of an *in vitro* formed SOD1 fibril showed two distinct fibril cores, one at the N-terminus of SOD1 (residues 3-55) and one at the C-terminus (residues 86-151) (66). In this model, I18 and I35 both contribute to the formation of two separate hydrophobic cavities, of which there are three, in the N-terminal SOD1 fibril core. Meanwhile, A95 contributes to the formation of the first C-terminal core, whereas I104 and I149 contribute to the second. Our data would suggest that the N-terminal amyloid core is the one potentially forming in disease, through the I18- and I35-associated cores. We must be careful however to consider that the C-terminal core is potentially inhibited from forming in our model, in which SOD1 is C-terminally tagged with AcGFP. However, we reason that the N-terminus of SOD1 is the key determinant of its aggregation for three reasons. Reason one is the existence of several C-terminal ALS-causing truncation mutations in SOD1, including a variant that results in the expression of a 35 residue N-terminal fragment (67, 68). Reason two is that previous investigation of the effects of N-terminal tagging of SOD1 with AcGFP (the tag we used in this study) showed that N-terminal tagging resulted in an inability of SOD1 mutants to aggregate (34). Reason three is that binary epitope mapping of SOD1 from *in vitro*, *in cyto*, and *in vivo* models consistently show that the N-terminus is inaccessible to antibodies, whereas the C-terminus is variably reactive depending on model and mutation (17, 19, 23, 28, 69).

We observed that our 7P variant was highly unfolded and destabilised, yet was not rapidly degraded by cells. Rather, aggregation prone variants were more rapidly degraded by cells in our model, and the 7P variant was degraded at a rate similar to SOD1-WT. Previous work investigating the half-life of SOD1 variants has shown that mutant forms of SOD1 are more rapidly degraded in cells than SOD1-WT, with the ubiquitin proteasome being the major contributing pathway of degradation (46, 70). Blocking UPS-mediated degradation of our proline variants showed that I18P, I35P, and 7P were not affected by proteasome inhibition, suggesting that the cellular mechanisms that recognise misfolded SOD1 variants as substrates for the UPS may not recognise these ones. The exact mechanism(s) by which cells recognise misfolded SOD1 remains unclear, although molecular chaperones in the cytoplasm and endoplasmic reticulum have been implicated (71, 72). Our data suggest that the amyloidogenicity of ^4^VQGIINFE^21^ and ^30^KVWGSIKGL^38^ are necessary for UPS-mediated degradation of SOD1 since our I18P and I35P variants did not become more insoluble with UPS inhibition. We cannot determine exactly how this effect is taking place, whether through inhibition of SOD1 oligomer formation or whether these two sites interact with protein quality control machinery remains to be elucidated. Future research using proximity-based ligation mass spectrometry would be useful in determining the differences in the interactomes between our proline variants and ALS-associated SOD1 mutants, potentially identifying unique interactors (73).

Based on the present work, we propose several conclusions and future lines of research. We show strong evidence to suggest that amyloidogenic segments ^14^VQGIINFE^21^ and ^30^KVWGSIKGL^38^ are the key regions governing the intracellular aggregation and propagation of SOD1 aggregation. Furthermore, we show evidence that suggests the existence of a specific recognition mechanism of misfolded SOD1 in cells, rather than unfolded SOD1, as evidenced by the unstable 7P variant being degraded at a similar rate to stable WT SOD1. Finally, our evidence supports that ALS-associated mutations in SOD1 need not be inherently toxic, but rather may enhance already toxic properties present within the SOD1 protein that are typically obscured due to stability of the protein. Future investigations should include proteomic-based experiments to identify interactors of toxic misfolded SOD1 in comparison to unfolded SOD1 to reveal those interactors that are specific for toxic forms of SOD1. Investigations into the cooperativity between ^14^VQGIINFE^21^ and ^30^KVWGSIKGL^38^ should also be performed, including determining if a β-hairpin composed of β-II and β-III is responsible for misfolded SOD1 oligomer and aggregate genesis. Finally, mapping the entire SOD1 variant landscape using deep mutational scanning has the potential to reveal further regions within the protein that contribute to aggregation and toxicity, as well as provide information to more accurately curate clinically-relevant variants (74). Determining the above will aid in understanding the molecular pathogenesis of ALS and also in determining more effective treatments for this fatal neurodegenerative disorder.

## Methods

### In silico sequence analysis of Cu/Zn superoxide dismutase for amyloidogenic segments and mutations that disrupt them

The canonical wild-type SOD1 amino acid sequence (Uniprot:P00441), without the N-terminal methionine, was modified to incorporate the G85R mutation. Following this, the SOD1-G85R sequence was input into multiple sequence-based informatics tools for the determination of amyloidogenic sequence segments. These included; Aggrescan-2D (75), ZipperDB (32), MetAmyl (76), TANGO (77), WALTZ (78), FoldAmyloid (79), FISH Amyloid (80), Aggrescan-3D (1.0) (81), and Zyggregator (82). Importantly, for all analyses the default tool settings were used. In the case of Aggrescan-3D, chain A of the x-ray crystal structure PDB: 1HL5 (83) was used by inputting the G85R mutation and other proline mutations in the Aggrescan-3D tool. Amino acids that were identified as amyloidogenic according to the specific tool, had their frequency of identification summed to give the SOD1 amino acid index frequency of amyloid prediction. Identification of amino acids for proline substitution was performed by sequentially replacing amino acids in the identified amyloidogenic segments and choosing those that decreased the predicted amyloidogenicity to the greatest extent.

### Plasmids for Mammalian Protein Expression

The empty backbone plasmid pAcGFP-N1 (Addgene: #54705) was a kind gift from Professor Michael Davidson. C-terminally TdTomato-tagged SOD1-G85R has been previously described (29). C-terminally AcGFP-tagged SOD1 variants SOD1-WT (Addgene: #264074) and SOD1-G85R (Addgene: #26410) were kind gifts from Professor Elizabeth Fisher (University College London, United Kingdom) (34). Residues for proline substitution were identified as described above, and generated by site directed mutagenesis by Genscript (New Jersey, USA). Plasmids were heat transformed into subcloning efficiency chemically competent DH5α cells (Thermofisher, USA) and purified using endotoxin-free maxiprep kits (Thermofisher, USA) as per the manufacturer’s instructions.

### Mammalian Tissue Culture

U2OS cells (ATCC - HTB-96) were cultured in Advanced Dulbecco’s modified Eagles medium-F12 (Adv. DMEM-F12) (Invitrogen, USA), supplemented with 10% (v/v) heat inactivated fetal bovine serum (FBS) (ThermoFisher, USA) and 2 mM GlutaMAX (ThermoFisher, USA). NSC-34 cells(84) were cultured in Adv. DMEM-F12 with 10% FBS and 2 mM GlutaMAX (ThermoFisher, USA). HEK293-T cells were cultured in DMEM-F12 with high glucose and 10% FBS and 2 mM GlutaMAX. In order to passage and plate U2OS, HEK293-T, and NSC-34 cells, they were washed once with pre-warmed 1× PBS (ThermoFisher, USA) and treated with 0.25% trypsin, 0.02% EDTA dissociation reagent (Invitrogen, USA) to lift off the adherent cells. The cells were pelleted via centrifugation (300 × *g* for 5 min) and resuspended in pre-warmed culture media. Following washing, plates, cover slips and chamber-slides were seeded at a confluency of 40% and cultured at 37 °C in a humidified incubator with 5% atmospheric CO_2_ for 24 h prior to transfection (∼70-80% confluent). Cells were transfected with plasmid DNA (0.5 µg per well of a 24-well plate and 0.25 µg per chamber of an 8-well chamber slide unless stated otherwise) 24 h post-plating using Lipofectamine LTX (Invitrogen, USA) according to the manufacturer’s instructions. Cell lines were transfected using Lipofectamine LTX reagent according to the manufacturer’s instructions. Where cells were co-transfected, equal amounts of each plasmid (AcGFP and TdTomato) were used.

### Proteolysis of SOD1-AcGFP constructs

HEK293-T cells expressing SOD1-AcGFP constructs were lysed using ice-cold 1× PBS 0.5% Triton X-100 supplemented with Halt Protease inhibitor cocktail (ThermoFisher, USA) prior to being clarified via centrifugation at 20 000 × *g* for 20 min, after which the supernatant was taken and the pellet discarded. Supernatants had their concentrations determined via DC Assay (BioRad).

For proteinase-K proteolysis experiments, lysates were equalised to a concentration of 20 mg/mL total protein and equilibrated to room temperature for 30 min. A t = 0 timepoint was taken and mixed with SDS-PAGE sample buffer before addition of proteinase-K. Proteinase-K was added to lysate at a final concentration of 10 ng/mL and samples were taken at 2.5, 5, 15, 30, 60, 120, and 240 min post-addition. Proteinase-K was deactivated via mixing with SDS-PAGE sample buffer.

For fast parallel proteolysis, lysates were prepared as above but thermolysin was added to a final concentration of 1 µg/mL with the lysates on ice in PCR strips. Strips were capped and placed into a rtPCR thermocycler which was set to heat in gradient mode. Lysates were heated for 5 min prior to taking out the strips and adding SDS-PAGE sample buffer to deactivate the thermolysin.

### Immunoblotting

Samples were subject to SDS-PAGE using NuPAGE 4-12% Bis-Tris gels. Gels were electrophoresed for 1 h at 150 V. Following electrophoresis, protein was transferred to 0.2 µm PVDF membrane via wet transfer for 1 h at 100 V. Membranes were blocked in 5% skim-milk in TBST (Tris buffered saline with 0.1% Tween-20) for 1 h prior to incubation in primary antibody in TBST with 5% skim-milk (rabbit anti-GFP 1:10,000; Abcam - ab290) overnight. Membranes were washed 3 times (5 min) in TBST prior to addition of secondary antibody (goat anti-rabbit HRP 1:10,000; Dako - P00448). Blots were washed 3 times in TBST prior to development using SuperSignal West Pico Plus substrate (ThermoFisher, USA). Blots were imaged using a Gel Doc XR+ molecular imager (BioRad, USA).

### Cell fixation using paraformaldehyde

Cells were cultured for 48 h post transfection after which they were fixed as follows. To fix cells, they were washed once with pre-warmed 1× PBS prior to addition of prewarmed 4% PFA in PBS, in which they were incubated for 20 min. Following fixation, cells were washed twice in 1× PBS before imaging or storage at 4 °C in the dark. Stored cells were imaged within 48 h of fixation.

### Imaging of fixed cells to determine inclusions

Fixed cells were imaged using an Axio Observer.Z1 inverted microscope (Carl Zeiss AG, Germany) fitted with an A-Plan 10×/0.25 objective, LED light source, a 62 HE BFP/GFP/HcRED reflector, a 395-495-610 beam splitter, a Colibri LED light source, and AxioCam HighRes camera (Carl Zeiss AG, Germany). Hoescht-33342 was excited with Hoescht-33342 (40 ms exposure time - 50% light source intensity) was excited using 365 nm LED through a 350 – 390 nm filter and its emission was captured at 402 – 448 nm. AcGFP (50 ms exposure time - 40% light source intensity) was excited with a 475 nm LED through a 460 – 488 nm filter and its emission captured from 500 – 557 nm.

### Quantification of Inclusion Formation in Cultured Cells

We followed previously published methods of image processing to quantify inclusions in our cells (39, 40) with some changes. Briefly, we performed illumination variation correction on all images using the background method. Nuclei were identified using the *IdentifyPrimaryObjects* module and the cytoplasm was identified using the *IdentifySecondaryObjects* module. Following segmentation, overlays of images were checked for artefacts or incorrect segmentation and these images were removed from analysis (roughly 2% of images taken). Measurements to generate cytoprofiles were obtained using the *MeasureObjectIntensity, MeasureGranularity, MeasureObjectSizeShape* and *MeasureObjectiIntensityDistribution* modules. A user was blinded to the cell populations and trained a Random Forest Classifier algorithm to identify cells using at least 400 cells per bin (inclusion, no inclusion, dead cells) until the accuracy for each classification was above 95%. All images within a dataset were then scored. We quantified dead cells as a separate bin due to this class often showing within the inclusion category unless separated (dead cells were found to be classified primarily on their size being smaller and shape more rounded).

### AcGFP and TdTomato -tagged SOD1 Mutant Co-expression

Images were acquired using an inverted Axio Observer Z1 microscope (Carl Zeiss AG, Germany) equipped with an AxioCam HighRes camera (Carl Zeiss AG, Germany). A LD plan-Neofluar 20×/0.4NA Korr M27 objective was used to capture images with a 62 HE BFP/GFP/HcRED reflector, a 395-495-610 beam splitter, with a Colibri LED light source (Carl Zeiss AG, Germany). AcGFP (8 ms exposure time - 25% light source intensity) was excited with 460 – 488 nm light and its emission captured from 500 – 557 nm, and TdTomato (150 ms exposure time - 100% light source intensity) was excited with 568 – 608 nm light and its emission captured from 615 – 800 nm. Camera settings were 2× analogue gain with 1,1 binning mode acquiring 14 bit images at 1388 × 1040 pixel resolution in a 4 × 4 grid for adequate well coverage. Importantly, these image acquisition settings were optimized to minimize saturation of fluorescent signal so that the ratio of TdTomato and AcGFP within inclusions would be accurately determined. The microscope was controlled with Zen Blue 2.5 Pro software (Carl Zeiss AG, Germany). Analysis of the ratios of AcGFP to TdTomato within inclusions was carried out as previously described (12).

### SOD1-AcGFP Mutant Co-expression with mCherry-Ubiquitin

Images were acquired on an SP5 Confocal (Leica, Germany) equipped with PMT detectors (gain 600 V), an Argon laser source (20% power, 3% transmission), a DPSS 561 nm laser (3% transmission), and a 63×/1.2NA water immersion objective. Images were acquired in 512×512 pixel format, with frame-by-frame sequential scanning and 3 line averages. The pinhole was set to 2 airy units. Analysis of the ratios of AcGFP to mCherry within inclusions was carried out as previously described (12).

### Gel Filtration Chromatography of Cell Lysates

HEK-293 cells expressing SOD1-AcGFP variants were lysed and 200 µL of clarified PBS-extracted, DSS-treated cell lysates with ultracentrifugation and DSS-treated AcGFP were injected onto a Superdex 200 Increase (10/300GL) column (GE Healthcare), using the UltiMate 3000 UHPLC system (ThermoFisher, USA), and eluted at a flow rate of 0.5 mL/min with 1× PBS (pH 7.4). Absorption wavelengths for the UV detector were set at 280 nm and 475 nm.

### Time-lapse Live-cell Imaging of NSC-34 Cells to Determine Survival and Inclusion Formation

Time-lapse live-cell imaging was carried out on NSC-34 cells in 8-well microscope chambers 24 h post-transfection under humidified conditions with 5% CO_2_ at 37 °C. Images were acquired at 1388 × 1040 pixel resolution (3×3 grid) were acquired every 20 min using an A-Plan 10×/0.25NA objective mounted on an inverted Axio Observer Z1 microscope (Carl Zeiss AG, Germany) equipped with an AxioCam HighRes camera (Carl Zeiss AG, Germany) and a motorized stage. Live-cell imaging started 24 h post-transfection. AcGFP was excited similar to above methods with the exception that exposure time was 25 ms.

Live-cell image stacks were analysed by a blinded observer. Cells were tracked from the initial image (t = 0) and observed for death and/or inclusion formation. Those cells that survived the entire observation period were censored at 48 h.

### Fluorescence Recovery After Photo-Bleaching

U2OS cells were plated into 8-well chamber slides (µ-Slide 8 Well - Ibidi, Germany) co-transfected with SOD1-AcGFP and SOD1-G85R-TdTomato constructs and cultured for 24 h as described above. Cells were transferred to an inverted SP5 laser scanning confocal (Leica, Germany) with an incubated chamber box. Within the chamber, cells were maintained at 37 °C under humidified conditions at 5% atmospheric CO_2_. A 63×/1.2NA water immersion objective was used to image cells where AcGFP and TdTomato were excited with an argon laser at 488 nm (70% laser power and 2% transmission for imaging, 100% transmission for bleaching). The pinhole was set to 2 Airy units. A photomultiplier tube (PMT) detector was used to detect GFP (collection window 498 - 530 nm, PMT gain = 600 V) and TdTomato (collection window 590 - 650 nm, PMT gain = 600 V). Imaging settings were optimized for enhancing the speed of frame acquisition by using bidirectional scanning at a 128 × 128 pixel resolution with a digital zoom factor of 5 (400 nm pixel size). The FRAP wizard tool was used to perform FRAP by using the zoom in bleach function. The FRAP time series was composed of 10 pre-bleach frames (0.2 s per frame), 10 bleach frames (0.2 s per frame), 620 post-bleach frames (0.2 s per frame). The mean fluorescence intensity of the bleached ROI, background ROI, and whole-cell ROI were obtained using ImageJ (version 1.53)(85). The data were then uploaded to the easyFRAP web tool(86). Data were normalised using the ‘double normalisation’ method and curves were fit with a double exponential equation. Only those fits with an R^2^ above 0.8 were accepted for analysis of the mobile fraction and half-time of recovery.

## Author Contributions

**Conceptualization:** LM, NRC; **Methodology:** LM, NRC; **Validation:** LM, JN, CS, MS; **Formal analysis:** LM, JN, CS, MS; **Investigation:** LM, JN, CS, MS; **Resources:** NRC, SSP; **Data curation:** LM, JN; **Writing - original draft:** LM; **Writing - review & editing:** LM, JN, CS, MS, NF, JJY, SSP, NRC; **Visualization:** LM; **Supervision:** LM, SSP, NRC; **Project administration:** LM, NRC; **Funding acquisition:** LM, SSP, NRC.

## Acknowledgments

LM acknowledges funding from Motor Neuron Disease Australia and FightMND. SSP acknowledges funding from CIHR Transitional Operating Grant 2682, NRC acknowledges grant support from ALS-Canada, Brain Canada, the R. Howard Webster Foundation, and the Canadian Consortium for Neurodegeneration in Aging. NRC also gratefully acknowledges generous donations from John Tognetti and William Lambert. The authors acknowledge the facilities and technical staff of the Djavad Mowafaghian Centre for Brain Health. All authors gratefully acknowledge the late Professor Justin J. Yerbury (University of Wollongong, Australia) who fought ALS through research and on a personal level.

