## Supporting Information for "Amyloidogenic regions in beta-strands II and III modulate the aggregation and toxicity of SOD1 in living cells"

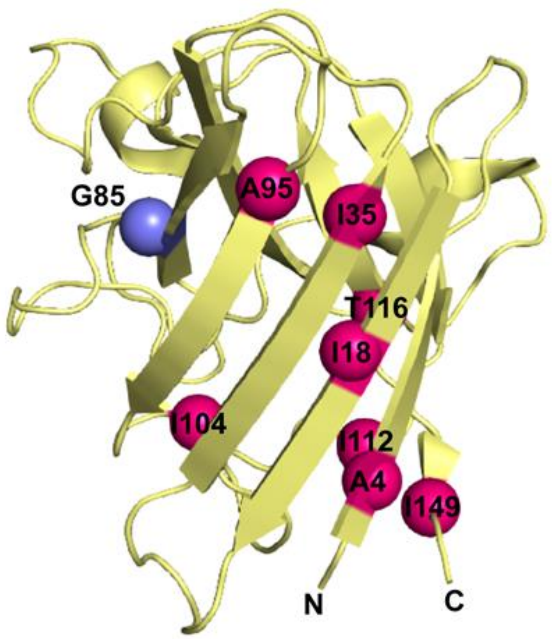


**Supplementary Figure 1. Location of all residues in SOD1-G85R (blue sphere) mutated to Proline (dark pink spheres) in this research. Crystal structure is 1HL5.**

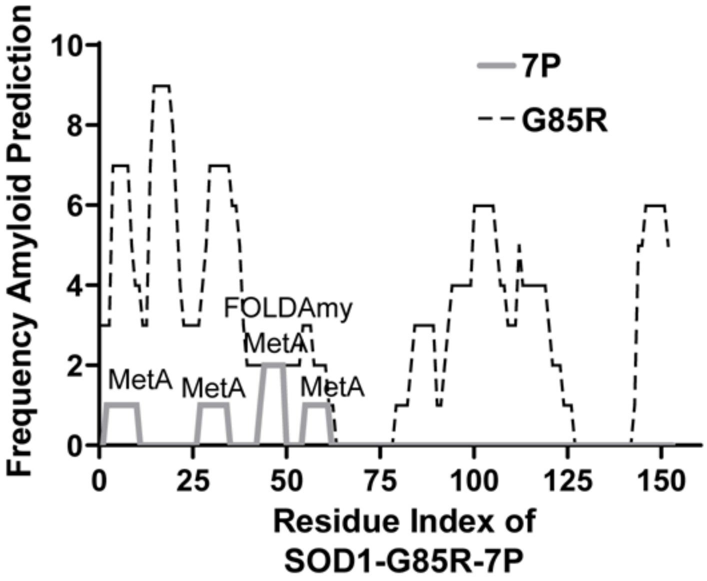


**Supplementary Figure 2. Prediction of amyloidogenic regions in the 7P (7 proline substitutions) variant of SOD1-G85R (grey) in comparison to SOD1-G85R (dashed lines). MetA and FOLDAmy show the tools that still predicted some amyloidogenicity from 7P.**

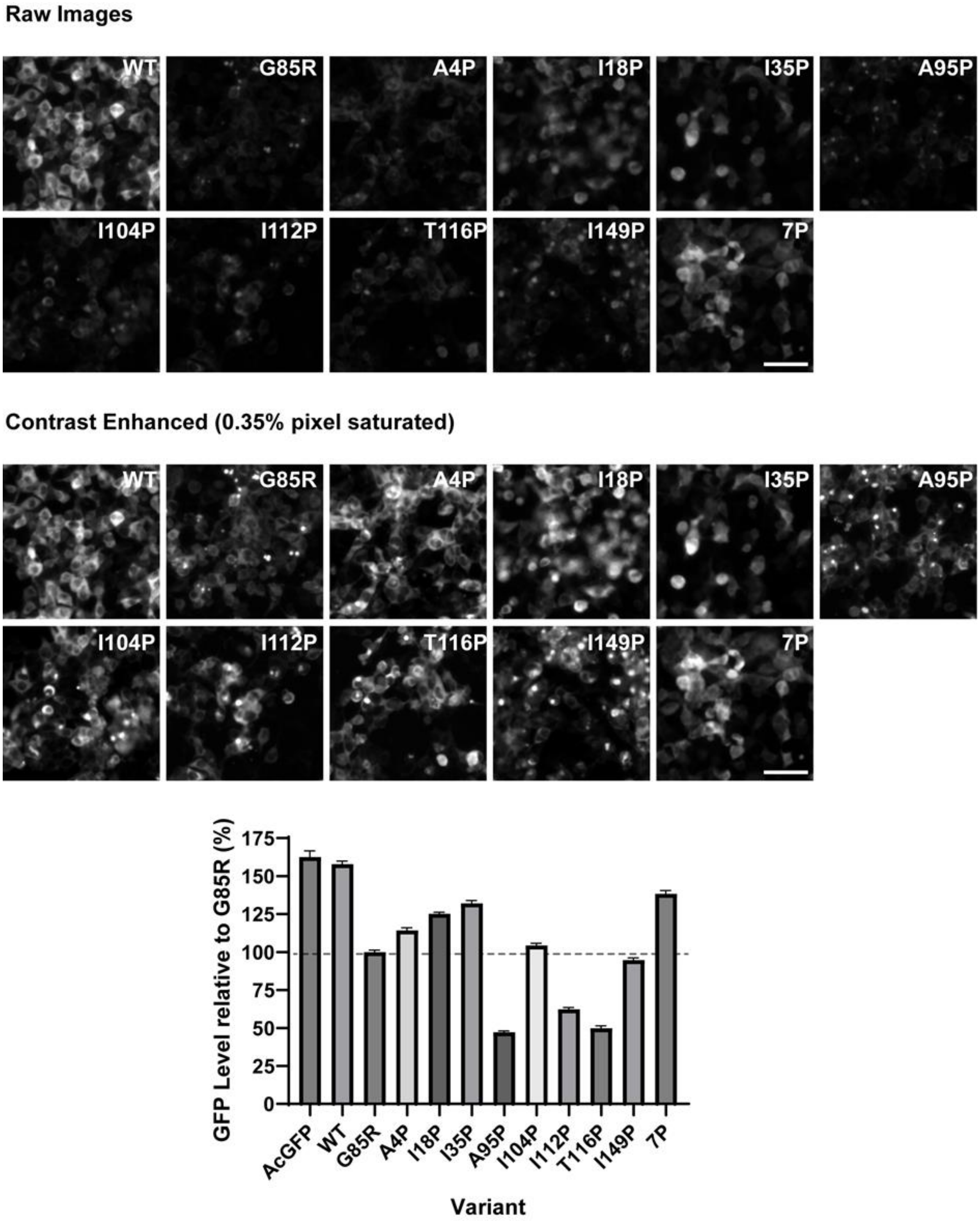


**Supplementary Figure 3. Mean levels of GFP fluorescent signal from HEK293T cells expressing each SOD1 variant normalised to SOD1-G85R.**

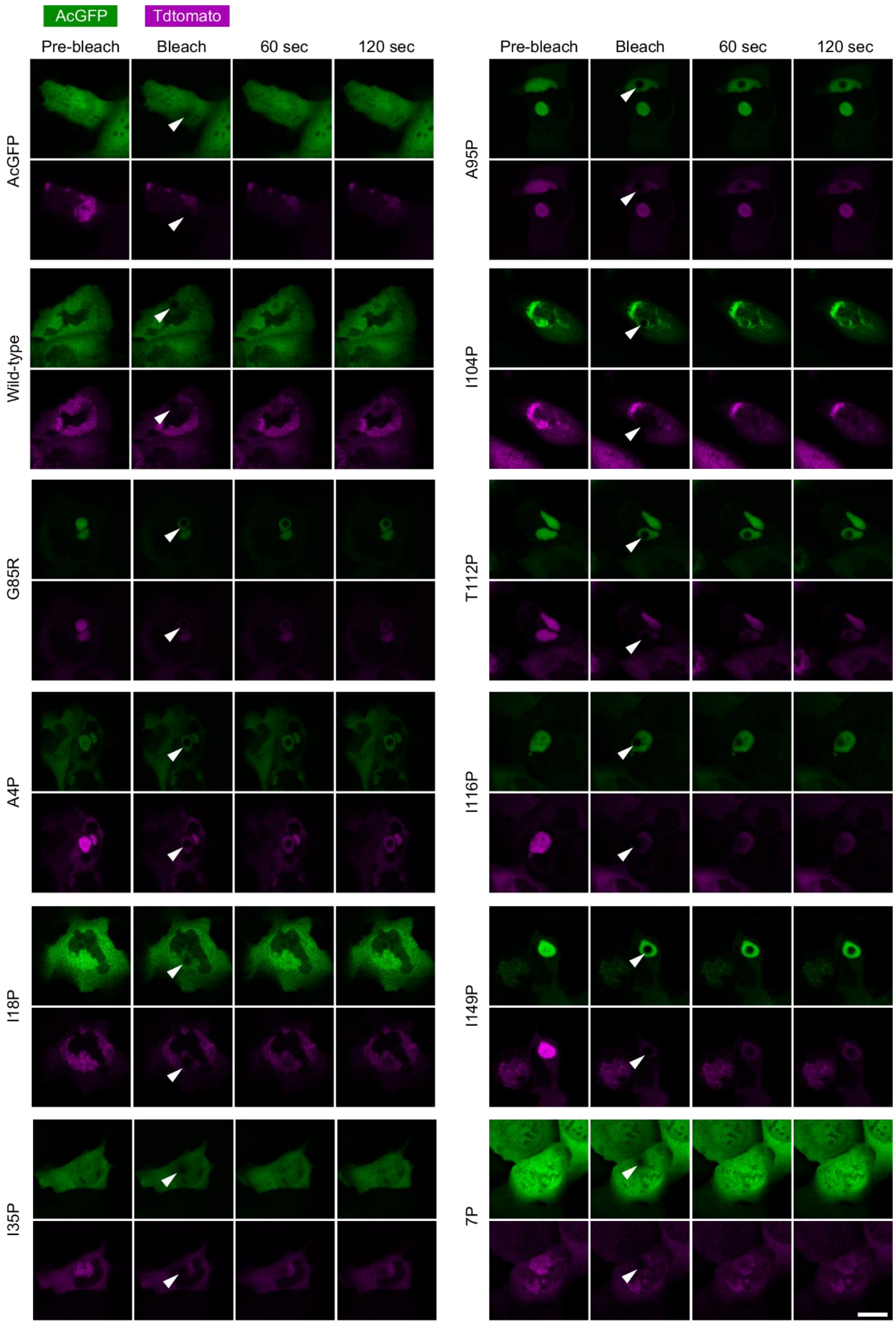


**Supplementary Figure 4. Examples of fluorescence recovery after photobleaching of transfected U2OS cells.**

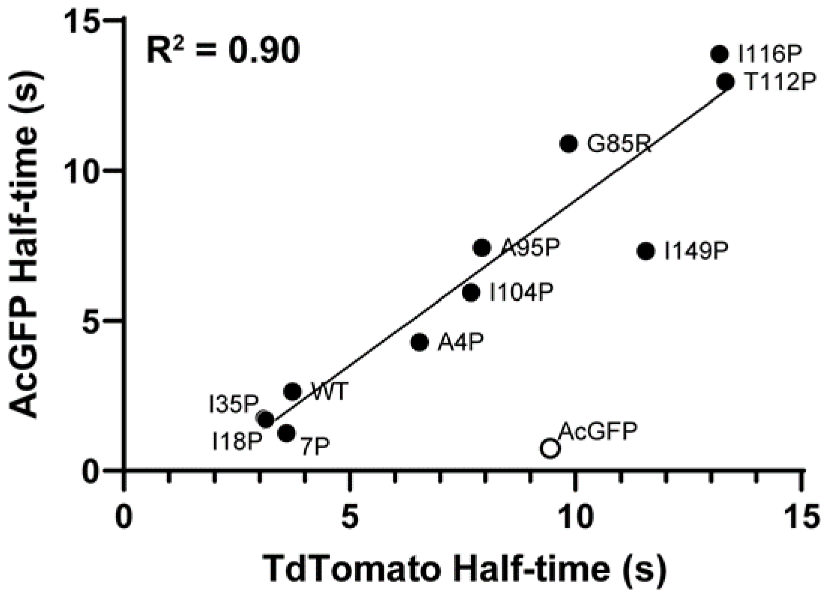


**Supplementary Figure 5. Correlation of TdTomato and AcGFP half-times to recovery. Unconjugated AcGFP (empty circle) was not included in the correlation.**

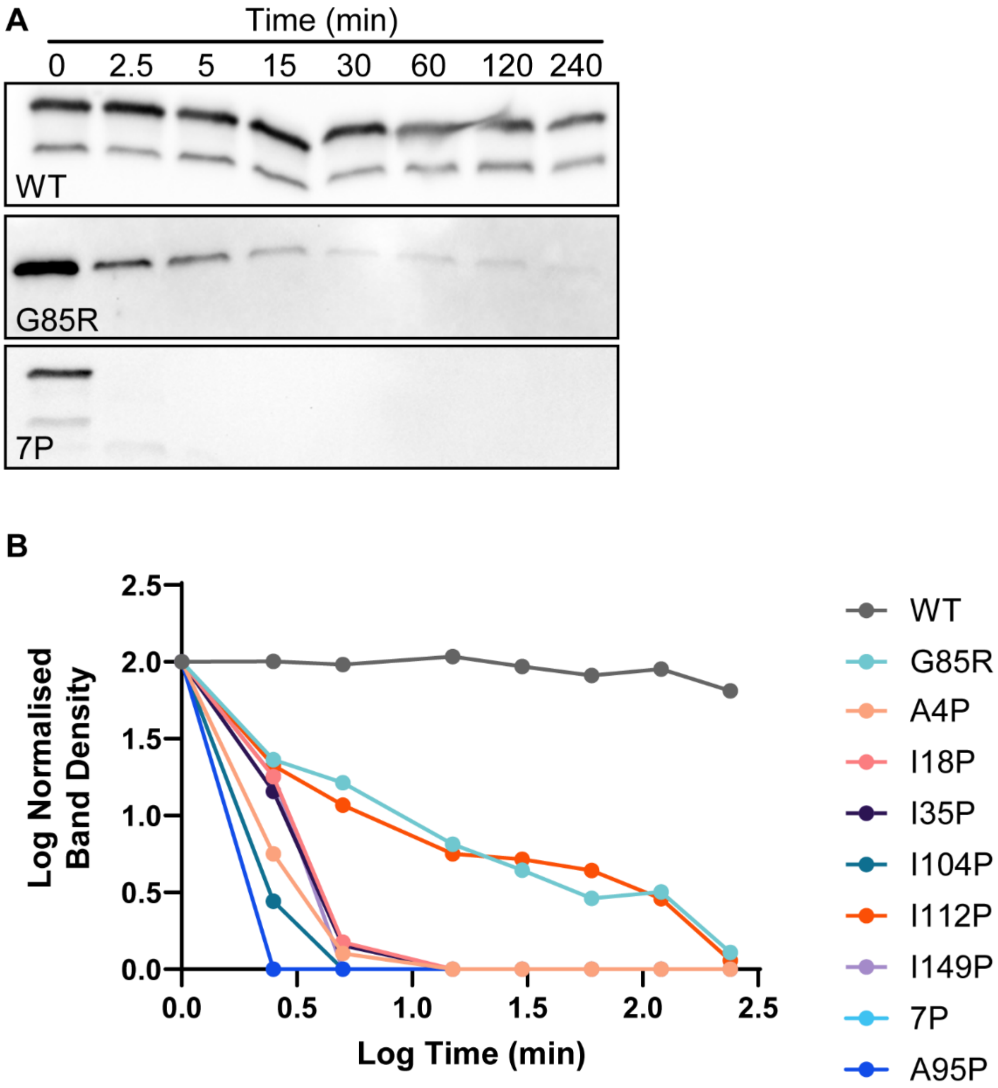


**Supplementary Figure 6. Proteinase-K digestion of SOD1 variants in lysates from transfected HEK-293T cells. (A) representative blots of WT, G85R and 7P showing protein signal disappearing with time. (B) plots of relative protein signal intensities compared to t = 0 band for each variant. Experiment is n = 1 replicate.**

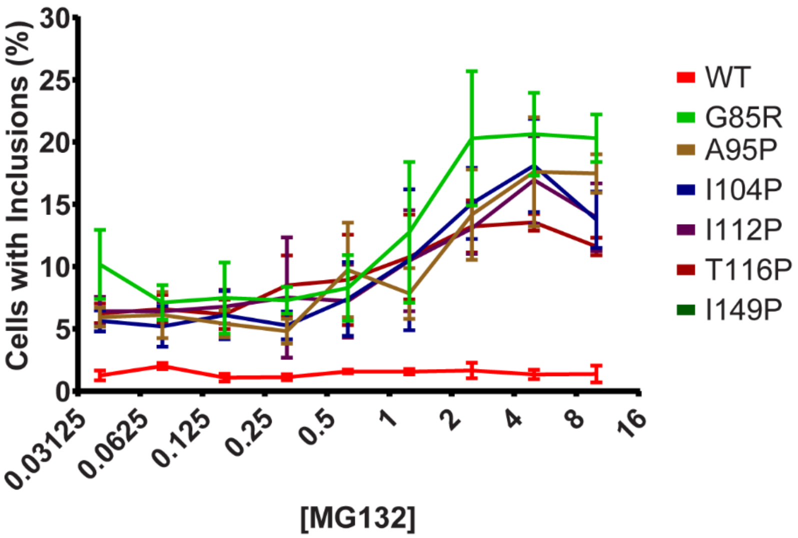


**Supplementary Figure 7. Comparisons of SOD1 proline variants to WT for MG132 (in μM) induced inclusion formation in HEK293T cells.**
